# Learning the pattern of epistasis linking genotype and phenotype in a protein

**DOI:** 10.1101/213835

**Authors:** Frank J. Poelwijk, Michael Socolich, Rama Ranganathan

## Abstract

Understanding the pattern of epistasis – the non-independence of mutations – is critical for relating genotype and phenotype in biological systems. However, the complexity of potential epistatic interactions has limited approaches to this problem at any level. To develop practical strategies, we carried out a comprehensive experimental study of epistasis between all mutations that link two phenotypically distinct variants of the *Entacmaea quadricolor* fluorescent protein. The data demonstrate significant high-order epistatic interactions between mutations, but also reveals extraordinary sparsity, enabling novel experimental strategies and sequence-based statistical methods for learning the relevant epistasis. The sequence space linking the parental fluorescent proteins is functionally connected through paths of single mutations; thus, high-order epistasis in proteins is consistent with evolution through stepwise variation and selection. This work initiates a path towards characterizing epistasis in proteins in general.

## Introduction

The central properties of proteins – folding, biochemical function, and evolvability – arise from a global pattern of cooperative energetic interactions between amino acid residues. Knowledge of this pattern is essential for understanding protein mechanism and evolution. However, the problem is extraordinarily complex. Energetic cooperativity in proteins is one manifestation of the more general principle of epistasis, the non-independent contributions of the parts that make up a biological system^1^. Epistasis can occur at the pairwise level (two-way) or extend to a series of higher-order terms (three-way, four-way, etc.) that describes the full extent of possible interactions^2–5^. As a consequence, the number of potential epistatic interactions grows exponentially with the number of positions in a protein, a combinatorial problem that becomes rapidly inaccessible to any scale of experimentation. Indeed, the theoretical complexity of this problem is such that it is not feasible or rational to propose an exhaustive mapping of epistasis for any protein.

How, then, can we practically characterize the architecture of epistatic interactions between amino acids? We reasoned that a strategy may emerge from a focused experimental case study in which we make all possible combinations of mutations within a limited set of positions within a protein – a dataset from which we can directly determine the extent of epistasis and explore possible simplifying methods. As a model system, we chose the *Entacmaea quadricolor* fluorescent protein eqFP611^6^, a protein in which spectral properties and brightness represent easily measured, quantitative phenotypes with a broad dynamic range. Recently, two variants of eqFP611 have been reported, one bright deep-red (mKate2, λ_ex_=590nm, λ_em_=635nm^7^) and one bright blue (mTagBFP2, λ_ex_=405nm, λ_em_=460nm^8^), that are separated by thirteen mutations (Fig 1A); we will refer to these as the “parental” genotypes.

**Figure 1:**
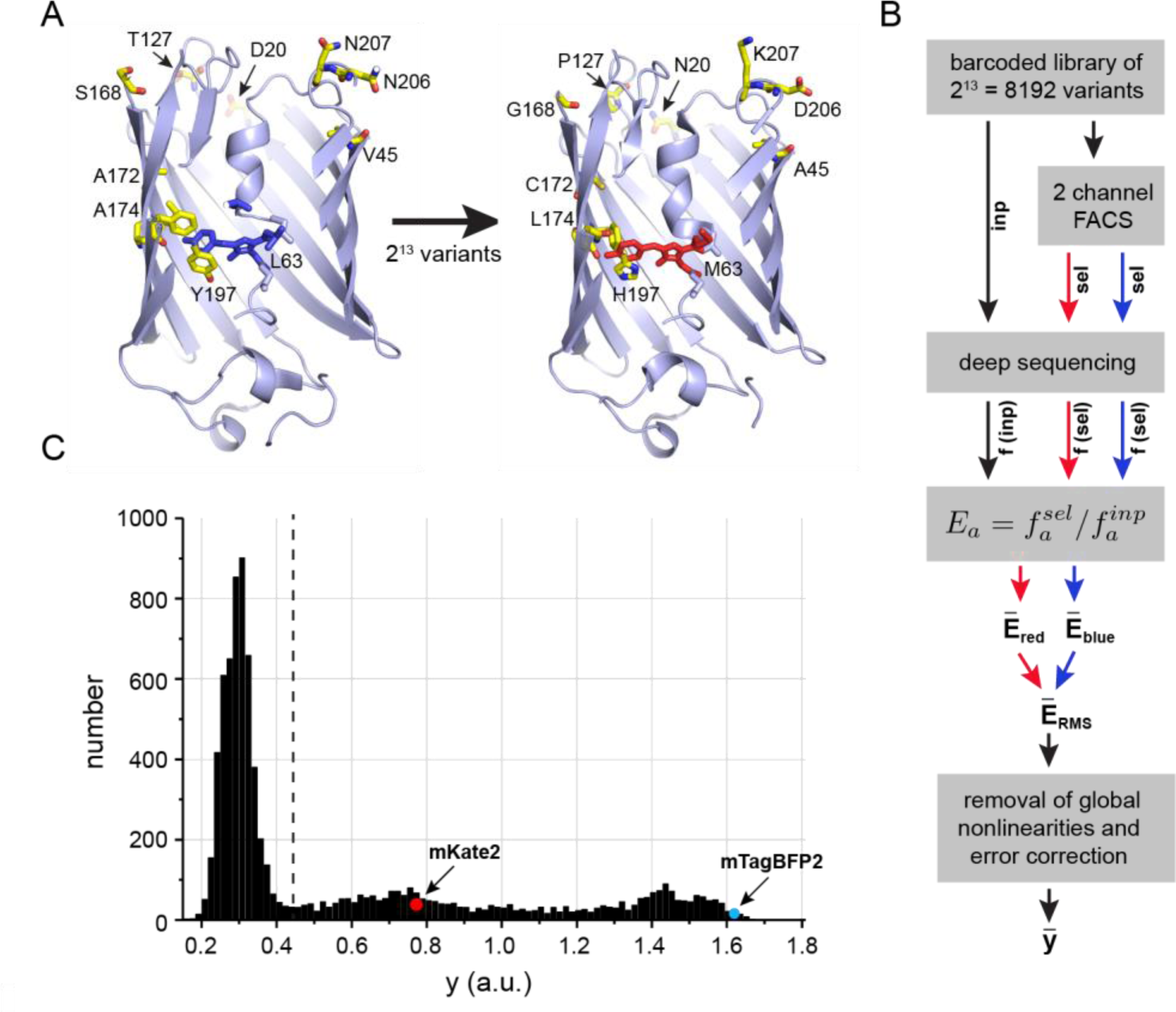
Combinatorial mutagenesis and data collection. **A**, mTagBF2 (left) and mKate2 (right) are blue and red variants, respectively. of the *Entacmaea quadricolor* fluorescent protein eqFP611 ^6^ that differ by 13 amino acid substitutions (10 shown). This defines a total sequence space linking the two of 2^13^=8192 variants. **B**, A schematic of the experimental protocol in which every variant is assigned a quantitative phenotype 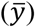 – brightness, a combination of fluorescence in both red and blue channels (see Methods for details). The phenotype is computed such that the independent action of mutations corresponds to additivity. C, The distribution of phenotypes for all 8,192 variants; the dashed line corresponds to the detection threshold for fluorescence.

By developing new technologies for high-throughput combinatorial mutagenesis and quantitative phenotyping, we explored the full space of 2^13^ = 8192 variants comprising the parental genotypes and all possible intermediates between them. These data reveal a broad range of high-order epistatic interactions between mutations, demonstrating non-trivial complexity in the relationship between genotype and phenotype. However, we find that epistasis is also highly sparse compared to theoretical limits, a property that opens up the use of powerful computational techniques for uncovering the epistatic architecture with limited experimental or sequence data. Despite the existence of high-order epistasis, we find that the blue and red variants of eqFP611 are fully connected through paths of single mutations in which protein function is maintained throughout. Thus, high-order epistasis is nevertheless consistent with phenotypic variation through stepwise variation and selection, a condition that should facilitate evolution.

## Results

### A complete combinatorial mapping of phenotypes

We developed an efficient iterative gene synthesis approach to simultaneously construct and barcode the full library of 8192 fluorescent protein (FP) variants that represents the parental genotypes mKate2^7^ and mTagBFP2^8^ and all possible intermediates (Fig. 1B, Extended Data Fig. 1, and Methods). This strategy makes it possible to readout the identity of every combination of mutations simply by high-throughput DNA sequencing of the barcode region. We expressed the library in *Escherichia coli*, carried out two-color fluorescence activated cell sorting (FACS) to select variants by brightness, and deep sequenced the input and selected libraries (Fig. 1B). The brightness of every FP allele *a* is given by its frequency *f*_*a*_ in the selected population relative to the input population: 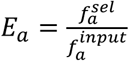. The color channels are normalized by the measured spectral properties of the parental red and blue proteins and are combined to produce a single quantitative phenotype – brightness – that is used in this study (Fig. 1B-C). Brightness integrates various underlying biophysical properties – extinction coefficient, quantum yield, or protein expression – and is therefore a rich phenotype for characterizing epistatic effects of mutations.

Trivial global nonlinearities in the dataset are expected just due to the experimental process and must generally be removed in assessing true epistatic interactions between mutations. To do this, we used the procedure of linear-nonlinear optimization^9^ to apply a simple polynomial transform (*y* = *E*^0.44^), which minimizes global non-linearities (Extended Data Fig. 2 shows robustness of conclusions to this process). The result is a quantitative assignment of phenotypes for all 8192 variants in a form in which independence of mutational effects corresponds to additivity in *y*. The data show that the parental genotypes are brightly fluorescent but many of the intermediates are not (Fig. 1C) – a first indication that we can expect substantial epistasis between mutations linking the two.

### From phenotypes to epistasis

From the full dataset of phenotypes, we computed the complete hierarchy – 1-way, 2-way, 3-way, 4-way, etc. – of epistatic interactions between the thirteen mutated positions.

Mathematically, epistasis is a transform (**Ω**) from the space of phenotypes 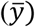 of individual variants to a space of context-dependent effects of the underlying mutations (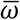, Fig. 2A)^2^:

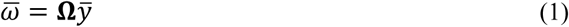

**Figure 2:**
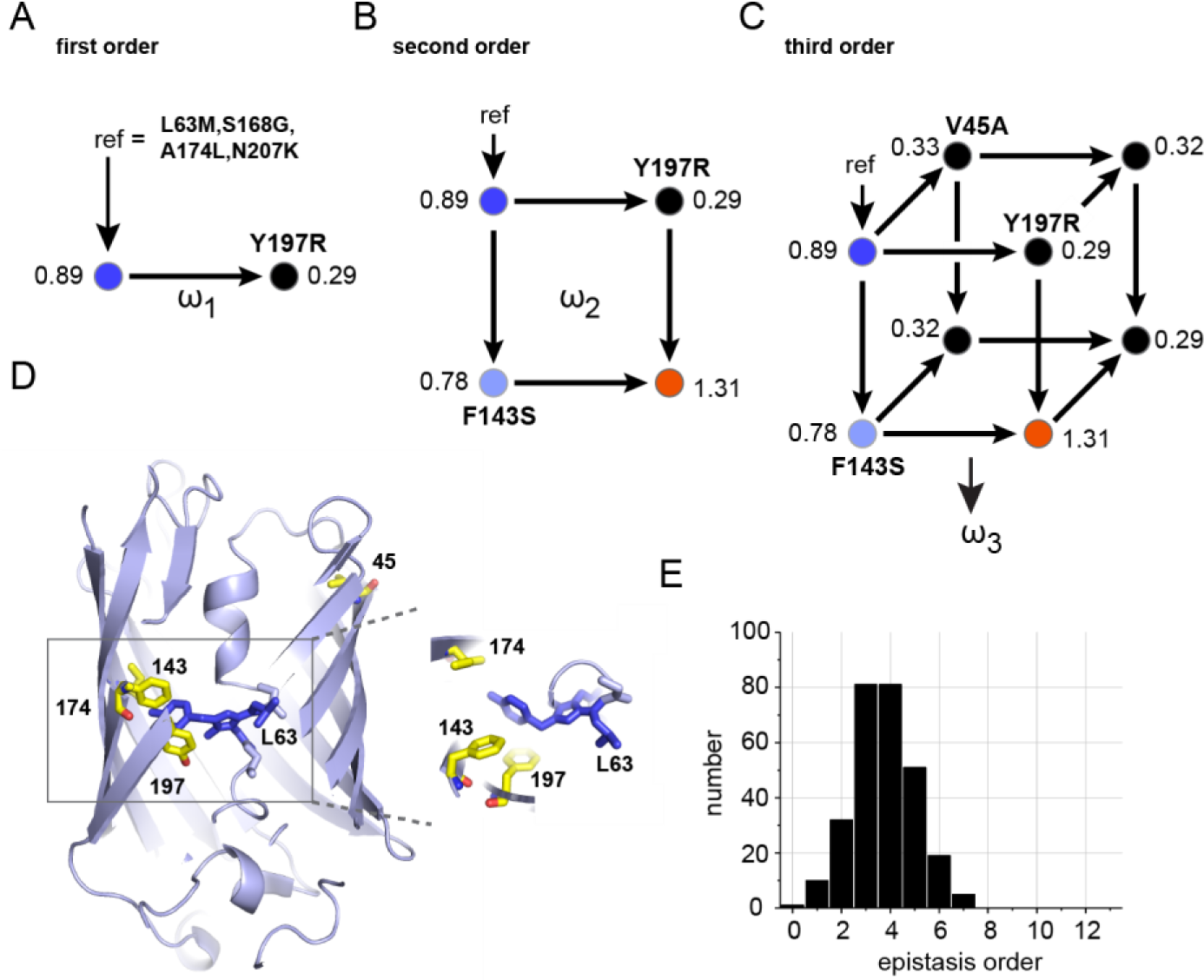
High-order epistasis in the sequence variations linking mTagBFP2 and mKate2. For illustration, panels A-C show the single reference form of epistasis taking an arbitrary genotype as the background (ref = L63M/S168G/A174L/N207K); circles indicate fluorescence color. All other panels indicate background-averaged epistasis. **A**, First-order epistasis is simply the effect of a single mutation. For example, in the reference background, Y197R shows *ω*_1_ = 0.29 − 0.89 = −0.67, indicating loss-of-function. **B**, Second-order epistasis is the dependence of a first order term on a second mutation. Here, Y197R has a completely different effect in the background of F143S, and thus these two mutations display a large second-order term (*ω*_2_ = 1.13). **C**, Third-order epistasis is the dependence of a second-order term on a third mutation. Here, the pairwise epistasis of Y197R and F143S is quenched in the background of V45A, indicating a large third-order term (*ω*_3_ = −1.15). **D**, Positions involved in large epistatic interactions are shown, indicating sites both proximal and distal to the chromophore. **E**, The distribution of the 280 statistically significant background-averaged epistatic terms (threshold, *p* < 0.01), showing a broad range of high-order interactions between amino acids, including terms up to the seventh order.

For *N* positions with a single substitution at each position, *N* is a vector of 2^*N*^ phenotypic measurements in binary order (here, 2^13^) and 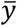 is a vector of 2*N* corresponding epistatic interactions. A first-order epistatic term (*ω*_1_) is the phenotypic effect of a single mutation, a second-order epistatic term (*ω*_2_) is the degree to which a single mutation effect is different in the background of second mutation, and a third-order epistasis (*ω*_3_) is the degree to which the second order epistasis is different in the background of a third mutation. Higher-order terms follow the same principle, such that an *n*^*th*^ order epistatic term is the degree to which an *n* − 1 order term depends on the context of yet another mutation, comprising a hierarchy of possible couplings between mutations. A key point is that 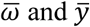 contain exactly the same information, but simply differ in its organization; 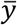 represents the phenotypes of individual variants while 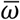 represents the non-additive interactions between the mutations.

A few examples help to explain the concept of epistasis. If we take, for illustration, the variant L63M/S168G/A174L/N207K as an arbitrary reference state (*y*_ref_ = 0.89, blue fluorescence) the data show that introducing the mutation Y197R results in reduced brightness (*y* = 0.29) (Fig. 2A). The difference in these two values defines a first-order epistasis (*ω*_1_ = *y*_*Y197R*_ − *y*_ref_ = −0.6). However, in the background of F143S, the effect of Y197R is entirely different; it shows increased brightness (*ω*_*1|F143S*_ = +0.53), with conversion to red fluorescence. This indicates a large second-order epistatic term (*ω*_2_ = *ω*_1|F14*S*_ − *ω*_1_ = 1.13, Fig. 2B), meaning that the effect of Y197R is context-dependent on F143S. This second-order term is itself dependent on other mutations. For example, in the background of V45A, the second-order epistasis between Y197R and F143S nearly vanishes (*ω*_2|*V45A*_ = −0.02), indicating a large third-order epistasis (*ω*_3_ = *ω*_*2|V45A*_ − *ω*_2_ = −1.15, Fig. 2C). These findings show that Y197R, F143S, and V45A work as a cooperative unit whose contribution to phenotype cannot be broken down into a simple, additive contribution of the underlying mutational effects. Instead, prediction of phenotypes involving these mutations requires knowledge of their individual effects and epistatic interactions at all orders.

In the examples discussed above, the effects of mutations are computed relative to a single reference genotype – the background in which the mutations are made. But, why should we restrict the definition of epistasis in the local mutational neighborhood of an arbitrarily chosen reference sequence? A more general analysis would be to compute each epistatic term as an average over all possible genetic backgrounds. For example, the effect of Y197R (the first-order epistasis *ω*_1_) can be computed not just with respect to a single reference (Fig. 2C), but as the average of its phenotypic effect in the background of every one of the other 2^*N−1*^ genotypes. Similarly, one can define background averaged pairwise, 3-way, and higher-order epistatic terms in which each term is averaged over all remaining genotypes. This view of epistasis is a global one, indicating the contribution of amino acids and interactions to protein function averaged over the full sequence space of variants^2,3,10–12^. We describe the profound distinction of single-reference and background averaged epistasis below.

We computed the background-averaged epistasis for the dataset of brightness phenotypes (see Methods). Analysis of error propagation provides a rigorous basis for establishing the statistical significance of epistatic terms as a function of order (Extended Data Fig. 3 and Methods). At a significance threshold of *P* < 0.01, we identify 280 background averaged epistatic terms, including many high-order interactions up to the 7^th^ order within the set of 13 mutated positions (Fig. 2E). Structurally, the epistatic terms involve not just residues in the local environment of the chromophore, but also includes positions (e.g. 45) located at a considerable distance at the opposite edge of the β-barrel (Fig. 2D). Indeed, the V45A mutation has the remarkable property of displaying a small effect on its own (*ω*_1_ = −0.09), but having much larger epistatic effects, for example, in modulating the pairwise coupling between Y197R and F143S (*ω*_3_ = −0.3). Overall, the data indicate a substantial number of high-order epistatic interactions between amino acid mutations that link the blue and red variants of the eqFP611 protein.

### Sparsity and reconstruction of phenotypes

The finding of significant epistatic terms up to the seventh order would seem to pose an insurmountable challenge to the goal of empirically relating genotype to phenotype in proteins. No studies are likely to make such measurements in general, and the scale of experimentation grows unmanageably with protein size. However, the data also suggest the possibility of extreme sparsity in the number of significant epistatic terms, a finding that if confirmed, can open up practical approaches. For example, the 280 significant epistatic terms identified here represent only a small fraction of the 8192 possible terms. How much information is encoded in just these terms? To study this, we used the inverse of the operation described in Eq. 1 to reconstruct the phenotypic measurements 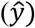 from any selected subset of background-averaged epistatic terms 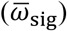:

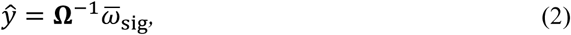

For 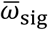 comprising all 280 statistically significant epistatic terms, a comparison of reconstructed phenotypes 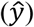 with actual phenotypes 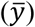 shows a goodness of fit coefficient (*R*^2^) of 0.98 (Fig. 3A and Methods), demonstrating nearly perfect agreement. This finding means (1) that the experiments performed in this work in fact correctly identify the major epistatic terms and (2) that nearly lossless data compression can be achieved (compression ratio 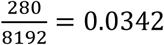) with the experimentally resolvable epistatic terms. The compression ratio serves as a quantitative measure of information content – the fraction of epistatic terms that suffice to predict phenotype up to a specified accuracy.

**Figure 3:**
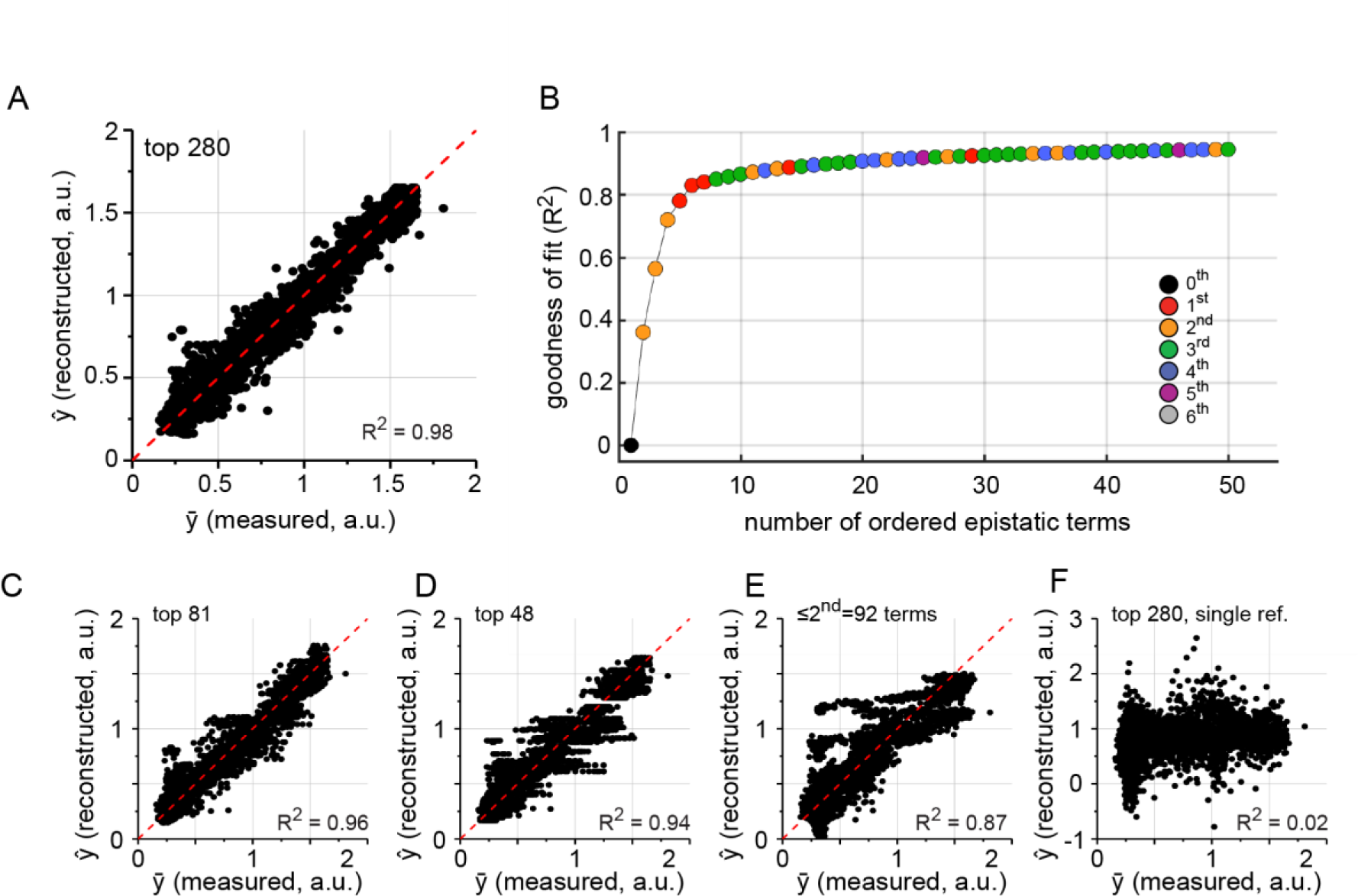
Sparsity in background-averaged (but not single-reference) epistasis. **A**, reconstruction of all phenotypes from the 280 significant background averaged epistatic terms displays excellent agreement with measured data (goodness-of-fit *R*^2^ = 0.98, see methods). **B**, A plot of goodness-of-fit against number of included epistatic terms arranged by degree of contribution, indicating extraordinary sparsity in information content. Colors show the order of epistasis. **C-D**, Consistent with sparsity, reconstruction of phenotypes with the top 81 (C) or top 48 (D) terms shows good agreement with measured data. **E**, Reconstruction with only second-order terms shows poorer agreement with data despite larger number of included terms, indicating the relevance of higher-order epistasis. **F**, Single-reference epistasis shows no predictive power in reconstructing phenotypes, indicating lack of sparsity in this form of epistasis. The plot shows the average of reconstruction for ten randomly chosen reference sequences.

To further examine the extent of sparsity, we calculated the goodness of fit between phenotypic data and prediction as a function of the number of included epistatic terms in 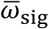, ordered by the size of their contribution to the explanatory power (Fig. 3B). The result demonstrates that information content in background-averaged epistasis is remarkably sparse; indeed, just the top 48 or 81 terms suffice to achieve an *R*^2^ = 0.94 or 0.96, respectively (Fig. 3C-D), suggesting information compression ratios of 0.005 or better can be achieved while still retaining accurate phenotype predictions. In contrast, retaining only epistatic terms up to the second order yields weaker predictive power (*R*^2^ = 0.87) despite a larger absolute number of terms (compare Figs.3C-D with 3E). Thus, phenotypes are optimally represented by a small number of epistatic terms, but these range from low to high order.

In contrast, sparse encoding of phenotypes is not evident when epistasis is not background-averaged (Fig. 3F). Indeed, with a particular genotype is taken as a reference for mutational effects, there is no predictive power of phenotypes globally (Fig. 3F, *R*^2^ = 0.02), and reasonable values for *R*^2^ are only achieved by inclusion of all terms up the 11^th^ order (Extended Data Fig. 4). This indicates essentially no information compression by this approach to epistasis; that is, epistasis is not sparse if it is not background averaged. It is interesting to note that the typical mutant cycle experiment in biochemistry represents an instance of single-reference epistasis in which mutations are seen as perturbations of a “wild-type” state.

Taken together, these data show that many high-order epistatic interactions exist among the mutations introduced, but that with background-averaging, the distribution of information is highly sparse, such that a small number of terms suffice to specify all phenotypes to good accuracy.

### Practical approaches for mapping epistasis

Background-averaged epistasis provides an efficient, low-dimensional representation of protein phenotype, but this observation does not provide a practical solution to identifying the relevant terms. By definition, background-averaging (at any epistatic order) requires complete knowledge of phenotypes for all combinations of mutants. So, how then can we practically deduce the relevant epistatic terms? One approach comes from the field of signal processing: the theory of compressed sensing (CS) states that if a signal displays sparsity in some representation, it is possible to accurately reconstruct the signal from a limited, well-defined number of measurements through an optimization procedure that enforces the sparsity in that representation^13^. For proteins, this implies that we should be able to learn the relevant epistatic architecture with just a sparse sampling of mutant phenotypes.

We made a simple implementation of the CS algorithm (by L1 norm minimization) and find excellent estimation of the top background averaged epistatic terms from a small fraction (~6-11%) of mutants (Fig. 4A-B). Using Eq. 2, these estimated terms can then be used to predict all protein phenotypes with excellent accuracy (Fig. 4C-D). A scan over the number of mutants used for prediction shows that the top epistatic terms are asymptotically well-estimated from very modest samplings of mutational variants (Fig. 4E). Note that this procedure does not simply amount to systematically sampling the low-order mutants; instead, the key is to sparsely sample over the space of all mutant combinations – a prescription for experiment design that is apparently best-suited for systems with sparse, high-order epistatic constraints. It will be important to understand how the sampling of mutations – the size of the experiment – scales with the number of sequence positions undergoing variation – the size of the problem. Since the latter grows exponentially, it seems likely that the degree of sparsity will be an even greater constraint for larger problems. Thus, these results provide a foundation for a rational experimental approach for learning the epistatic architecture of proteins.

**Figure 4:**
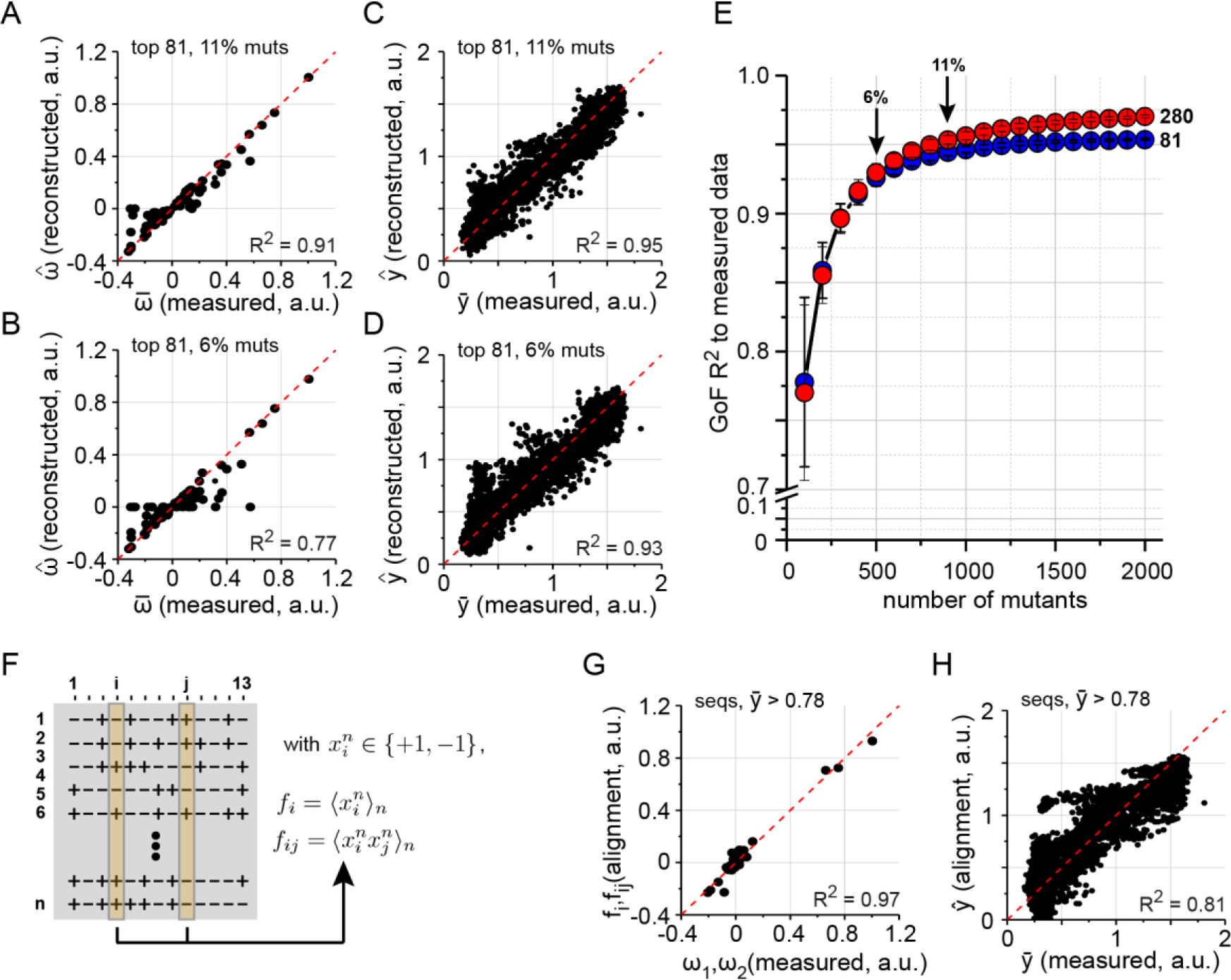
Practical strategies for learning the epistatic structure. **A-B**, Estimation of the top epistatic terms (including a broad range of high-order terms) from random samplings of phenotypes for 6-11% of variants, using the method of compressive sensing (CS). **C-D**, Reconstruction of all phenotypes from the estimated epistatic terms in panels A-B. The data show excellent approximation of relevant high-order epistasis and prediction of phenotypes from sparse sampling of data. **E**, Goodness of phenotype prediction for all 8,192 variants as a function of CS-based estimation of epistasis (top 81 or 280 terms) from many trials of sampling the indicated number of variants. **F**, A representation of a multiple sequence alignment (MSA) 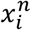 comprising *n* sequences by the *i* = (1 … 13) mutated positions; amino acids are represented by +1 and −1 to indicate residues in mTagBP2 or mKate2, respectively. From the MSA, we can compute the average value of each position 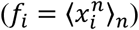, and the joint expectation of pairs of positions 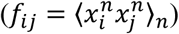, as indicated. **G-H**, Estimation of measured first and second order epistatic terms (G) and consequently, the ability to reconstruct all phenotypes (H) using only the first and second order alignment statistics.

A distinct approach is suggested by analyzing the statistics of amino acid frequencies in a sampling of functional sequences. For example, constructing a multiple sequence alignment of FP variants with brightness above the value of the red parent (*y* > 0.78) and representing the two amino acids *x* at each position *i* with −1 and +1 respectively, we find that the average value over all *n* sequences 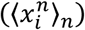 and the joint expectation between pairs of positions *i* and *j* 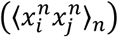 are closely related to the background averaged first-order and pairwise epistatic terms determined experimentally (Fig. 4F-G). This relationship holds even with sub-sampling of functional sequences included in the alignment (Extended Data Fig. 5). From these alignment-derived epistasis terms, it is possible to again quantitatively predict the phenotypes determined experimentally (Fig. 4H) to an extent that approaches what is possible with just limiting epistasis to the second order (Fig. 3E).

This result indicates a link between epistasis averaged over genetic backgrounds and statistical correlations averaged over sequences displaying function above a threshold, a condition analogous to the process of natural selection. This connection is the fundamental premise of coevolution-based methods that use amino acid correlations in multiple sequence alignments to estimate structural^14–16^ or functional^17–21^ couplings between residues in proteins. Though these methods have provided important insights^22–27^, our findings show that accurate phenotype prediction will require knowledge of higher-order epistatic terms as well. Such information is not formally included in current coevolution methods, but may be accessible if alignments are deep enough or the problem of epistasis in full proteins is sparse enough.

### Functional connectivity of the sequence space

How does the pattern of epistasis control the topology of the functional sequence space linking the blue and red variants of eqFP611? Indeed, the existence of severe forms of epistasis (e.g. sign epistasis or reciprocal sign epistasis^28^, in which intermediates can fall below the selection threshold) can limit and even abrogate the existence of single-step (or “connected”) paths between functional genotypes^29^. Thus, the study of the structure and connectivity of the space of functional genotypes is important for understanding how epistasis controls evolvability. In our dataset, nearly 50% of the statistically significant pairwise epistatic interactions represent cases of sign or reciprocal-sign epistasis (Extended Data Fig. 6), indicating that functional connectivity of the space linking the parental variants of eqFP611 is not trivial.

Figure 5A shows the network of all functionally connected 13-step paths – the “solution space” – between the blue and red parental variants (*y* > 0.78, the value of the red parent, and see Extended Data Fig. 7). The genotypes are colored according to fluorescence and edges represent single mutations between them. The data show (1) that the sequence space linking the parental genotypes is in fact fully connected at the functional threshold defined by these genotypes, (2) that solution space is shaped like a dumbbell, with two densities of functionally bright sequences near to the parental genotypes connected by a narrow neck, and (3) that the color switches at the neck. The shape of the network reflects the pattern of epistasis. For example, the narrowest part of the solution space occurs in the middle where the number of possible genotypes is the largest (Fig. 5B), indicating severe epistatic constraints on mutational paths at these steps with regard to retaining brightness along the path (Extended Data Fig. 7D).

**Figure 5:**
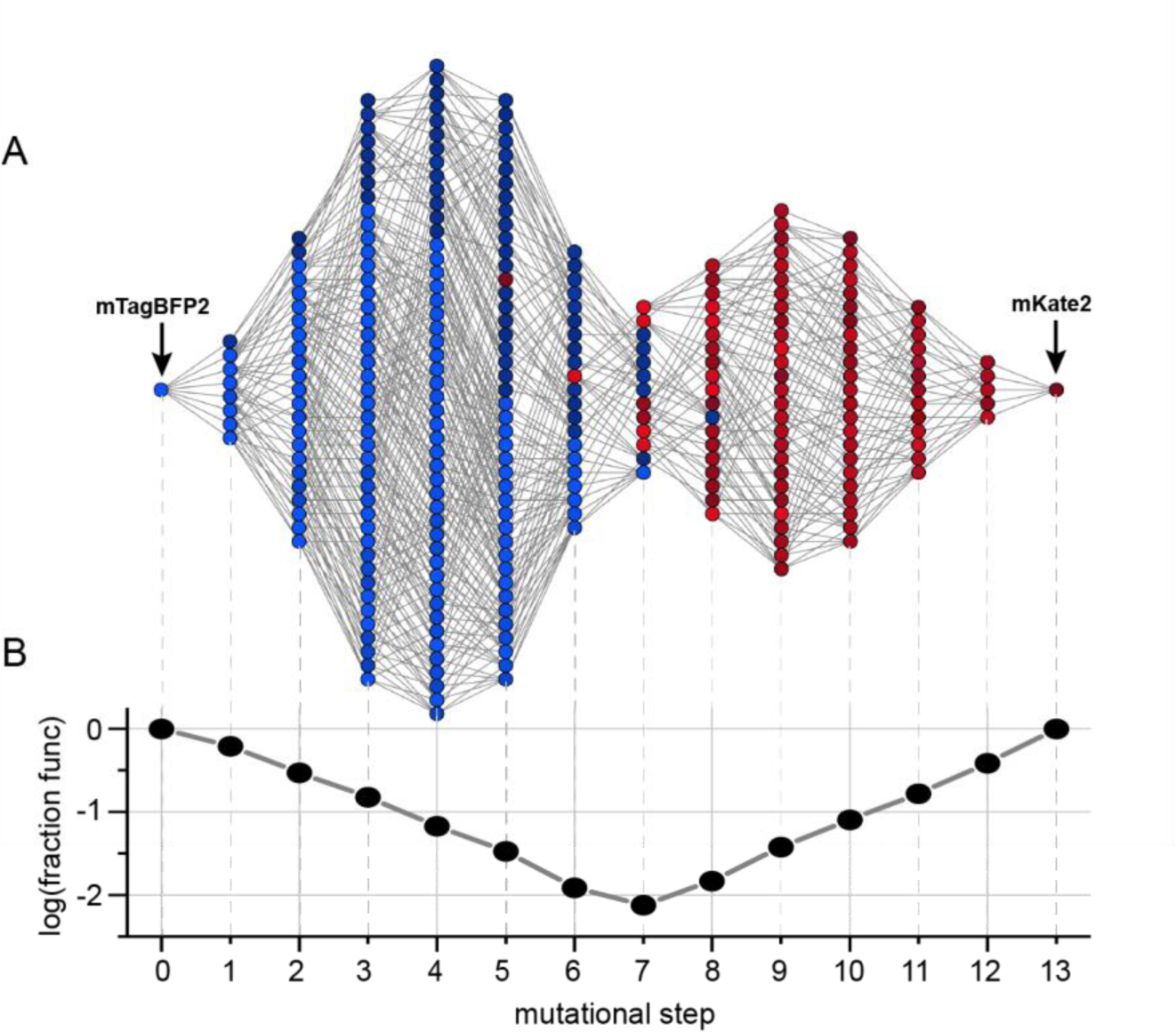
Single-step connectivity of the sequence space linking mTagBFP2 and mKate2. **A**, A graph of all genotypes comprising a set of sequences connected by single-step variation, as a function of mutational step from mTagBFP2 to mKate2. The brightness threshold for selection of genotypes is at the level of the mKate2. Thus, the sequence space linking the two parental genotypes is fully connected through single mutations without loss of parental function, and the shape of the solution space involves a thin neck near the middle. **B**, The fraction of connected genotypes at each step of mutation reinforces the notion that the space is most constrained at the thin neck, a consequence of severe epistatic constraints at the intermediate steps.

Though we focus on brightness as a phenotype in this work, it is also informative that the fluorescence color switches at the narrow neck. The blue and red spectral states arise from chemically distinct chromophores that are auto-catalytically generated upon folding from amino acids at positions 63-65^30,31^. Interestingly, the data show that L63M – the only chromophore mutation – is necessary but insufficient on its own to produce the red chromophore. Instead, red fluorescence requires several specifically-ordered mutational steps after L63M, another indication of epistasis in the path between the blue and red parental variants.

Overall, the connectivity of the solution space between the blue and red variants shows that the existence of high-order epistatic terms can nevertheless be consistent with evolution through stepwise variation and selection. The effect of epistasis is in specifying the topology of the solution space and in restricting the number of available paths. For example, at the specified brightness threshold, only 1.15 × 105 out of 6.23 × 109 paths between the blue and red parental genotypes (or, ~ 0.002%) are functionally connected. Single-step connectivity is an advantageous feature for phenotypic evolution through a process of random variation and selection. Thus, connectivity of the solution space represents a clear example of how constraints on protein sequences can arise from not just from the requirement to fold and function, but from the dynamics of the evolutionary process^25,32,33^.

### Conclusion

Defining the pattern of epistasis between amino acids is essential for understanding protein function and evolvability. Given the vast theoretical complexity of epistatic interactions between amino acids, it is essential to carry out model experimental studies as a basis for developing practical strategies. Here, we show significant high-order epistasis in the mutational landscape linking red and blue variants of the epFP611 fluorescent protein. But, with background-averaging, epistasis is also profoundly sparse, inspiring the use of powerful analytic tools for defining the epistatic architecture through sparse data collection. Conventional strategies for studying proteins focus on low-order mutagenesis^34–38^, but the data presented here suggest that a different experimental approach is optimal for defining the relationship between genotype and phenotype – limited sampling of all combinations of mutations, and sparse reconstruction to deduce the relevant epistatic terms.

Interestingly, second-order background-averaged epistatic terms are well approximated by the statistical correlations between amino acids in an alignment of functional protein sequences, providing support for yet another approach. Current alignment-based methods for deducing amino acid couplings in proteins either rely on analysis of conserved, collective correlations between positions^17,19^ or on inference of direct pairwise interactions through inverse methods in statistical physics^14,16,39^. The data presented here provide a critical benchmark for these approaches, defining the minimal epistatic terms that must be estimated in order to successfully relate genotype to phenotype. The sparsity of these epistatic terms may provide a productive constraint for developing a proper theoretical framework for using statistical coevolution to quantitatively predict protein phenotypes.

What controls the prevalence, distribution, and spatial architecture of epistasis in proteins? Why should it be sparse? One limit comes from physical considerations; for example, the forces that bind atoms mostly act locally in protein structures, a property that forces long-range epistatic terms to be built up from the coupling of local interactions. However, the finding that the blue and red variants of eqFP611 are connected by single-step mutations suggests the possibility of other constraints as well. For example, if evolution is facilitated by the stepwise functional connectivity of genotypes, then it is clear that any pattern of internal epistasis that is inconsistent with connectivity will be less fit, regardless of its own phenotypic value. The practical analysis of epistasis is the starting point for testing such ideas, and this work provides a foundation towards that goal.

## Acknowledgements

We thank O. Rivoire, M. Weigt, R. Monasson, S. Cocco, and V. Krishna for discussions, and members of the Ranganathan and Sander laboratories for critical review of the manuscript. We also thank the High Performance Computing Group (BioHPC) and the Genomics Core at UT Southwestern for providing computational resources and sequencing, respectively. This work was supported by NIH Grant RO1GM12345 (to R.R.), a Robert A. Welch Foundation Grant I-1366 (to R.R.), and the Green Center for Systems Biology at UT Southwestern Medical Center. F.J.P was an HHMI fellow of the Helen Hay Whitney Foundation.

## Author Contributions

F.J.P. and R.R. developed the research plan and experimental strategy. F.J.P. designed the strategies for library construction and carried out all steps of the experimental process shown in Fig. 1B. F.J.P. also designed the computational strategies for epistasis calculations and sparse reconstruction, and F.J.P. and R.R. carried out data analyses and simulations. M.A.S. contributed aspects of molecular biology, protein biochemistry, and spectroscopy. F.J.P. and R.R. wrote the paper.

## Methods

### Combinatorial library construction

The combinatorial library of 2^13^ FP variants was constructed by an iterative synthesis protocol in which mutant combinations and an associated barcode are co-assembled in a derivative of the pRD007 plasmid^40^ (Extended Data Fig. 1). Briefly, 34 DNA segments were synthesized (500bp gBlocks, IDT Inc), each comprising a portion of the FP coding sequence, a 5’ barcode encoding the identity of mutations within this region, and three restriction sites in between (a Type II site flanked by two non-pallindromic Type IIs sites). Barcodes are designed to have Hamming distance of at least two between each other, with each segment barcode comprising three bases plus a parity base (which represents the numeric sum of the three bases modulo four). The Type IIs sites permit scarless in-frame joining of segments by cutting outside of the recognition sequence and the Type II sites increases cloning efficiency by elimination of uncut or back-ligated species. Each segment encodes one to three mutated positions, with the most 5’ segment of the FP gene including an IPTG-inducible promoter from pTrc99A^41^ and a random 12bp “uniqueness” barcode that uniquely labels each individual clone. The FP genes are constructed iteratively 3’ to 5’, where at each step, one segment is ligated into the host vector, transformed into *Escherichia coli* DH5*α*^42^, grown overnight, and the resulting plasmid library isolated to serve the host vector for the next round. In this process, combinations of mutants and associated segment barcodes are assembled together. A key technique is the alternating use of two sets of type IIS and Type II restriction endonucleases (Extended Data Fig. 1). After complete assembly, the library is transformed into *E. coli* MC1061 (^43^, AVB100, Avidity Inc), at low DNA concentration (5ng DNA total) to suppress multiple transformants (typical library size of 5⋅10^6^). Sequences were optimized to avoid AGA and AGG codons, which are rare in *E. coli*.

### Cell sorting

MC1061 cells containing the FP library were grown at 37°C to an optical density of 0.8 in LB plus 50μg/ml kanamycin, induced with 200μM IPTG for one hour, and kept at 16°C overnight. Cells are then washed and resuspended in deionized sterile water, diluted to ~ 10^7^/ml, and sorted on a BD FACSAria (UT Southwestern Medical Center cytometry core) at excitation/emission wavelengths of 405/460nm and 532/610nm. The total sorted population was 6.2 x 10^7^ cells, and gating thresholds were chosen to recover the top 1% of cells in each channel. Gating by threshold still yields a graded output because single cells encoding any particular allele exhibit fluorescence that follows a near log-normal count distribution (see e.g. ^36^). Screening of colonies on plates yielded no evidence for multi-colored proteins, justifying stringent sorting thresholds along the observed phenotypic axes. Sorted cells were recovered in LB medium without antibiotics, grown overnight in LB plus 50 ¼g/ml kanamycin, and subject to plasmid isolation for deep sequencing.

### High-throughput sequencing, error correction, and phenotype determination

Samples for sequencing were prepared by PCR from plasmid libraries before or after selection using primers that incorporate Illumina adaptor sequences, a bar code specifying origin (input or output color channel), and a random stretch of five nucleotides in the initial 5’ region for phasing and cluster definition. Products were pooled in a ratio of 10:1:2 representing input, red, and blue output channels, and paired-end PE-100 sequencing was performed on a single lane of an Illumina Genome Analyzer IIx (UT Southwestern genomics core). Raw FASTQ files from the Illumina base-caller were processed with custom scripts in UNIX and MATLAB, and subjected to stringent quality filtering involving three criteria: 100% correct reads within a mask around the bar codes, correct specific barcodes, and a Q-score of at least 30 for each nucleotide in the random “uniqueness” barcode. Primer sequences and scripts are available upon request.

The uniqueness barcode provides a mechanism to correct for mis-sorting events and unobserved spurious mutations that can introduce errors in assigning phenotypes. For each allele-specific barcode *a* we compute input and output counts for each uniqueness barcode *k* as 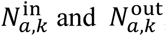, and a linear allelic enrichment 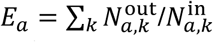. Modeling experimental errors as a Poisson-process, we compute noise on the output counts of a uniqueness barcode as 〈noise〉_*i*_ ∝ 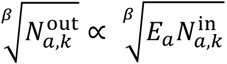, calculate the Z scores for the individual uniqueness bars as

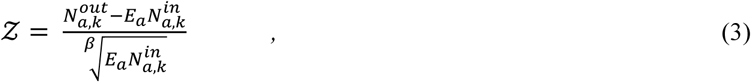

and set the upper and lower boundaries for inclusion in the data per uniqueness barcode as 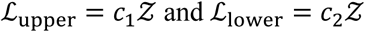. Choices of parameters (*β* = 1/0.35, *c*_1_ = 35, and *c*_2_ = 15) were based on robustness across color channels and for alleles over the full range of enrichments *E*_*a*_. Three rounds of outlier rejection led to removal of 2% of counts, after which final enrichments were calculated, and normalized by the known brightness of the red and the blue parental genotypes ^7,8^. MATLAB scripts are available upon request.

Analysis of epistasis requires elimination of trivial global nonlinearities in the data that arise from the experimental or analytic process. The general principle is that trivial nonlinearities will systematically influence every variant, while nonlinearities due to intramolecular epistasis are highly specific properties of a few variants. A logical approach is to find the simplest empirical transform 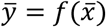 that minimizes the global non-linearity, especially in the most well determined (i.e. low-order) terms ^9^. We minimize 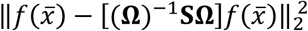, where **Ω** is the epistasis operator and **S** is a matrix which selects only epistatic terms up to order two. This led to 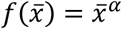 with *α* = 0.4364. Upon transformation, the data corresponding to genotypes with zero brightness are regularized by adding pseudocounts based on fitting noise present in non-functional genotypes. Extended Data Fig. 2 shows that the conclusions in this work are robust to these steps.

### Analysis of epistasis

For a single amino acid substitution at each of *N* positions, the full space of possible genotypes corresponds to 2^*N*^ individual variants. With phenotypes for all variants 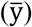 in a form that independence corresponds to additivity, the analysis of epistasis corresponds to a linear mapping 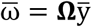, where **Ω** is a weighted Walsh-Hadamard transform, a class of generalized Fourier transforms. With background averaging, **Ω** = **VH**, where **V** and **H** can be recursively defined:

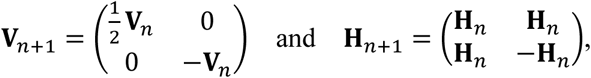

with **V**_0_ = **H**_0_ = 1, and *n* = {0 … *N* − 1}. For standard, single-reference epistasis, **Ω** = **VX**^***T***^**H**, where

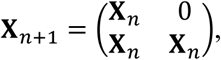

with **X**_0_ = 1. Conceptually, standard epistasis represents a local (Taylor) approximation of the fitness landscape expanded around one reference genotype, while background-averaged epistasis approximates the landscape in terms of its global features over the space of all possible genotypes. See ref^2^ for definitions and explanation.

### Functional trajectories and genotypic connectivity

To obtain the number of functional single-step trajectories, we compute an adjacency matrix between functional genotypes by binarization of the full genotype adjacency matrix above a select threshold brightness (for Fig. 5, *y* = 0.78, the value for the red parental variant). From the binarized adjacency matrix, the (*i, j*)-th element of the *m*^th^ power of the matrix gives the number of functional *m*-step trajectories that exist between genotypes *i* and *j*. Summing over the powers of this matrix to any order *M* gives all viable trajectories consisting of *M* or fewer steps in the sequence space. Conversion of the resulting summed matrix to block-diagonal form produces a “genotypic connectogram” – a graph that directly reveals the connectivity and topology of viable genotypes (Extended Data Fig. 7C).

### Sparse optimization and phenotype reconstruction

L1-norm minimization to find the optimal sparse distribution of epistatic terms was performed in MATLAB using the YALL1 solver, version 1.4 ^44^. The performance for mutant prediction is scored by the Goodness of Prediction (GoP) parameter 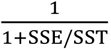, where SSE is the sum of squared errors between reconstruction phenotypes and the measured values, and SST is the total sum of squares. MATLAB scripts are available upon request.

---

**Extended Data Figure 1:**
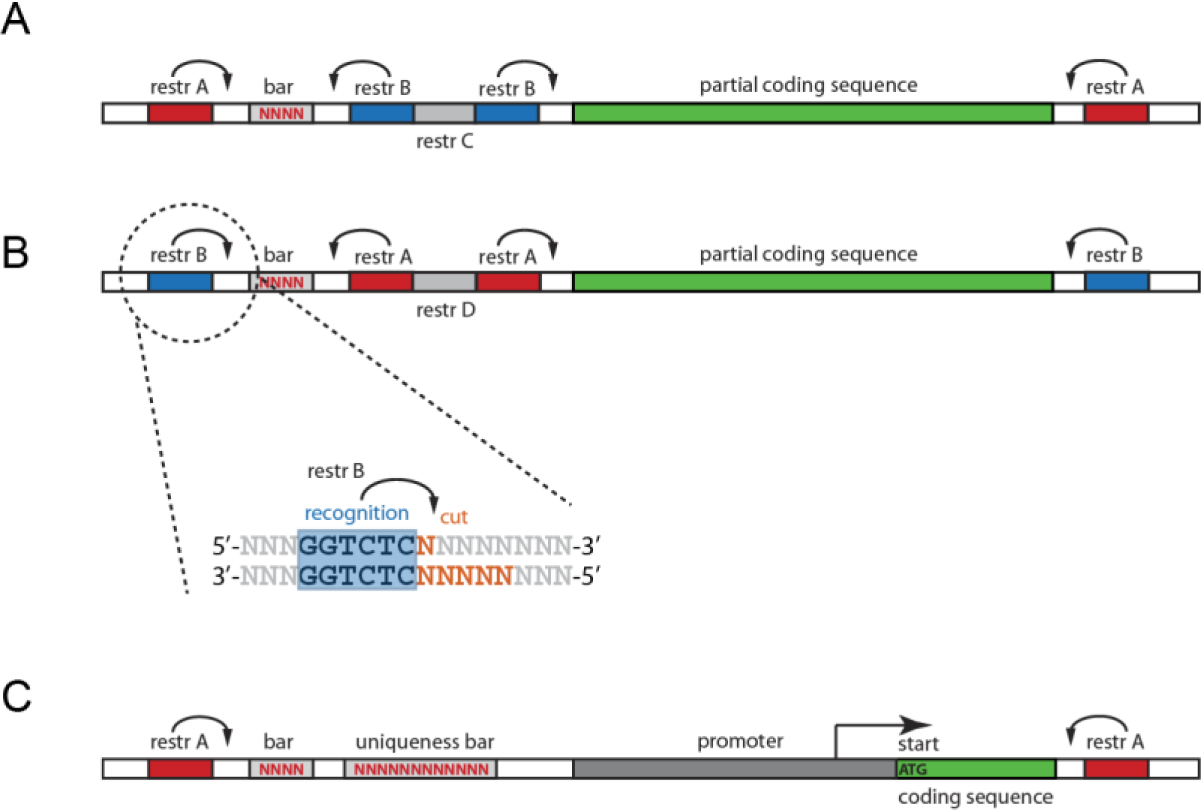
Combinatorial library synthesis. The library of FP variants is generated by sequential restriction and ligation steps incorporating pre-synthesized DNA segments (gBlock, IDT Inc.) from 3’ to 5’ that each contain a subset of the mutated positions and a partial barcode that indicates each combination of mutations. The number of segments is determined by a trade-off between minimizing the number of assembly steps and minimizing cost; here 34 construction segments were used – segments 1-4 with three mutations each (**A-B**) and a final fifth segment with one mutation and the promoter (**C**). Segments 1-4 have a design that alternates between the schemes shown in panels A and B, explained below. The barcodes are designed to have minimal Hamming distance of two between each other, and are embedded in a flanking sequence that is designed to avoid palindromes, long repeats, or restriction sites used in construction. The gene assembly process is as follows: The first construction segment is cut with Type IIS restriction enzyme BsrDI (restr A), and ligated to target vector pFPH, a derivative of pRD007 ^40^. After transformation and isolation of plasmid DNA, the resulting population of plasmids and the second segment are cut together with Type IIS enzyme BsaI (restr B), purified and ligated, inserting the second segment 5’ to the first and juxtaposing the segement barcodes. A “kill cut” is made with PstI (restr C) to reduce propagation of uncut or back-ligated species, and the ligation reaction is transformed and DNA isolated. This procedure is repeated for the remaining segments, alternating the use of the restriction enzymes as per panels A and B (restr D is NdeI). The final step incorporates the random “uniqueness” barcode.

**Extended Data Figure 2:**
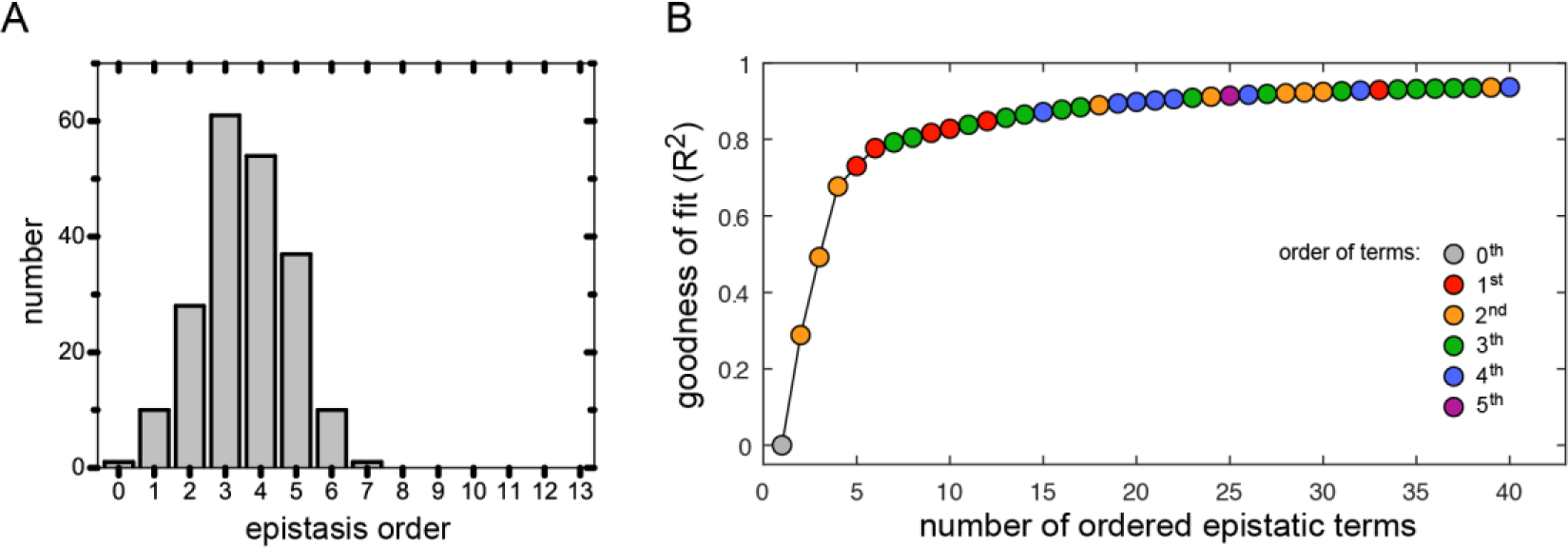
Epistasis analysis without removal of global non-linearities. **A**, Histogram of epistatic terms as a function of order at a significance threshold of *p* < .01 (after Bonferroni-Šidák correction for multiple testing), computed exactly as in Fig. 2G, but without the linear-nonlinear transform to minimize global non-linearity in the data. **B**, the Goodness-of-fit between measured and reconstructed phenotypes as a function of number of included epistatic terms, ordered by degree of contribution (analogous to Fig. 3B). The analysis shows that the basic conclusions of this work are, in this case, not strongly dependent on the nature of the phenotype transform.

**Extended Data Figure 3:**
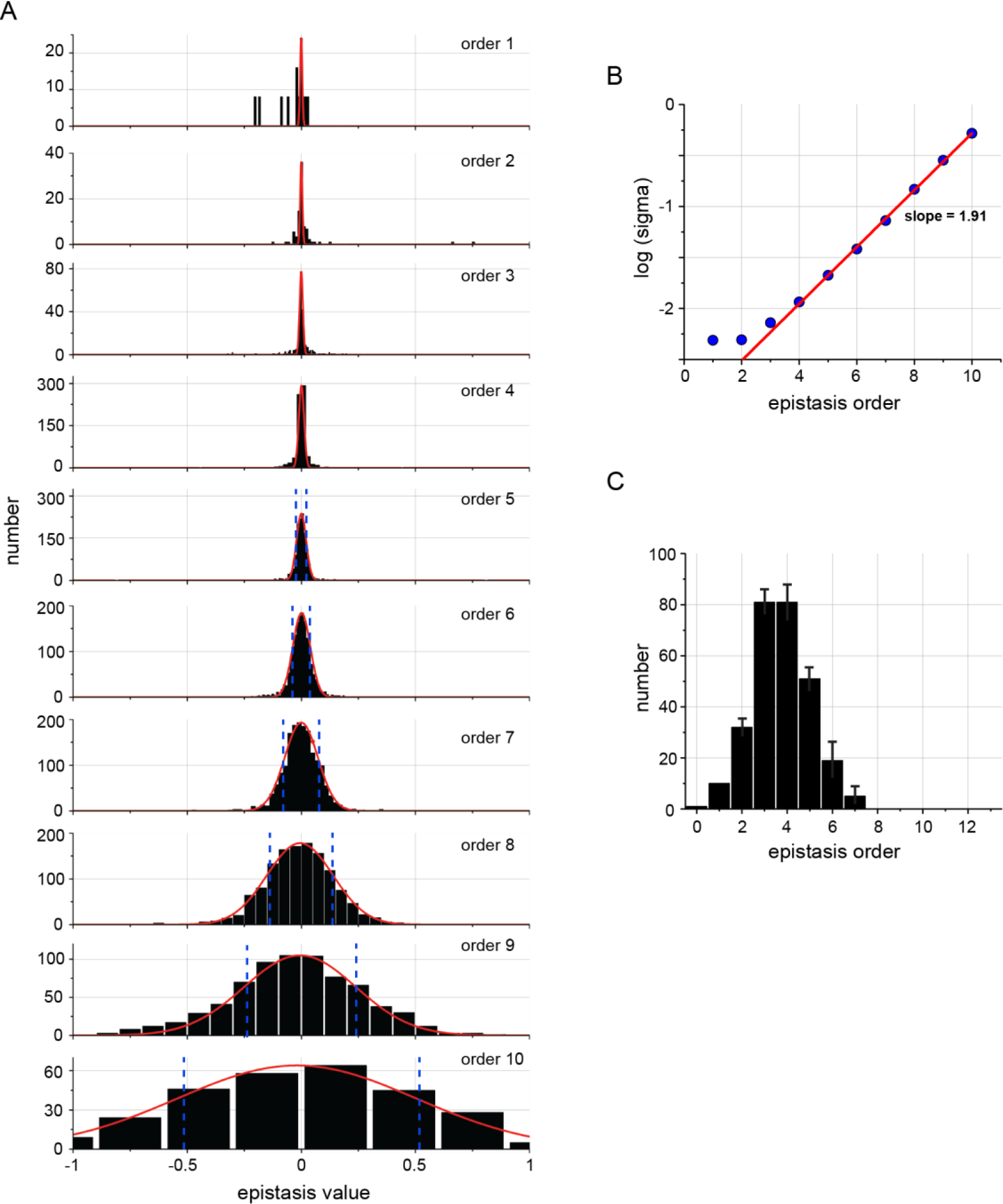
Error propagation and determination of significant epistatic terms. **A**, Histograms of calculated epistatic terms for orders 1-10, fitted to Gaussian distributions (in red). The data show that as expected by the propagation of error rule, the variance in the distribution grows with epistatic order. **B**, the logarithm of the fitted Gaussian widths of the histograms as a function of epistatic order. The slope indicates an increase in observed noise of a factor 1.91 per order, close to the theoretical expectation of 2 (*r*^2^ = 0.99). The y-axis intercept is at 6.1 x 10-4, suggesting an average per datapoint error of 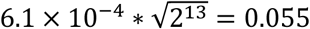. C, Number of epistatic terms as a function of order at a significance threshold of *p* = 0.01, after Bonferroni-Šidák correction (as in Fig. 2G). Here, “error bars” indicate the robustness of this distribution to the choice of p-value. The low-range is *p* = 0.001, and the high-range is *p* = 0.05.

**Extended Data Figure 4:**
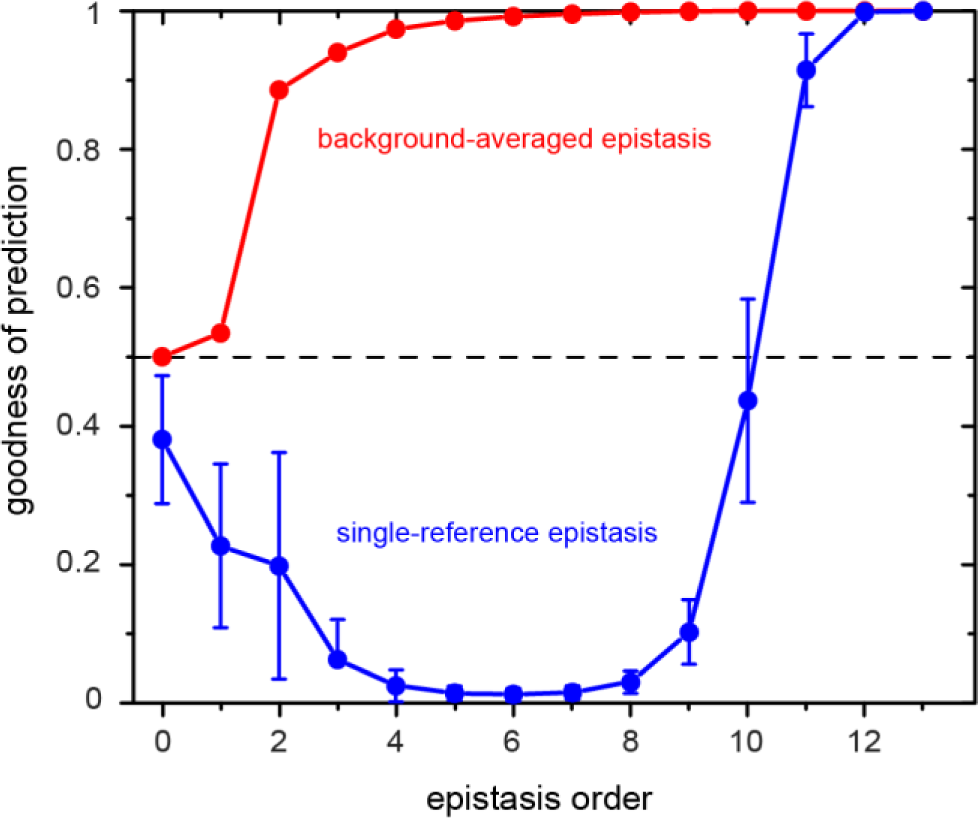
Error propagation and determination of significant epistatic terms. Phenotype prediction, a comparison of background-averaged and single-reference epistasis. The goodness of prediction (GoP, see methods) for epistatic terms computed using background averaging (red) or with taking a single genotype as a reference (blue, shown is the mean and standard deviation for 100 randomly chosen genotypes). A goodness of prediction of 0.5 is expected for a random (fully uninformed) prediction. The data show that prediction using single-reference epistasis only out-performs a uniformed prediction when terms up to the 11^th^ order are included (note that the GoP necessarily converges to unity when all orders of epistasis are included. Thus, epistasis is not sparse when using single-reference definitions.

**Extended Data Figure 5:**
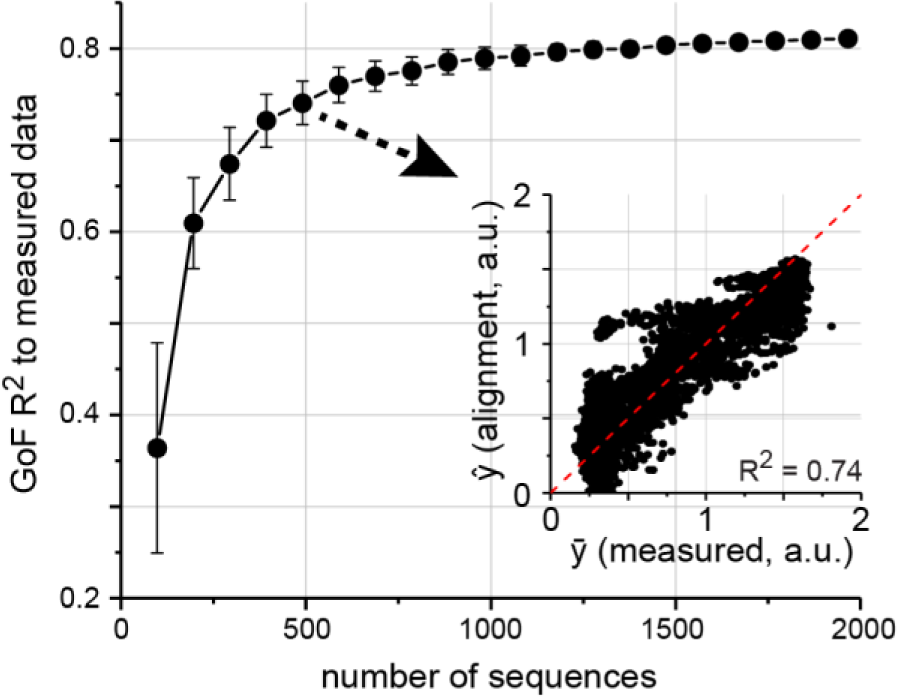
Phenotype prediction from alignment statistics, as a function of sequence sampling. The graph shows the goodness-of-fit R^2^ between measured and reconstructed data (as in Fig. 4H) for epistatic terms estimated from alignments of functional sequences sampled from the full alignment of functional sequences (defined as those with *y* > 0.78). The data show rapid convergence of phenotype prediction with even sub-sampling of functional sequences. The inset shows the quality of phenotype reconstruction for one instance of sampling 500 sequences from the full set of functional genotypes (compare with Fig. 4H).

**Extended Data Figure 6:**
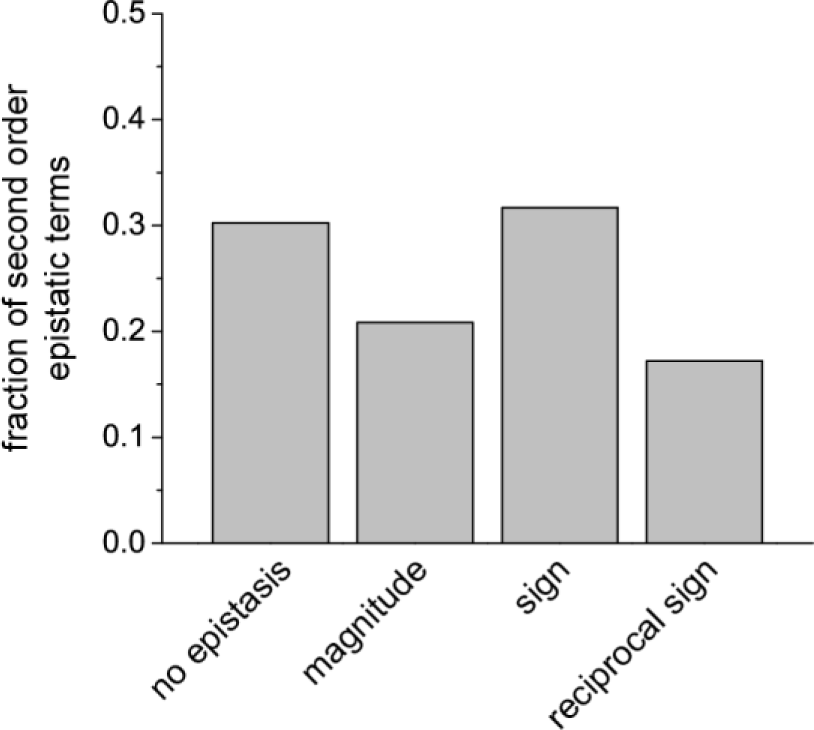
Distribution of epistasis type amongst the pairwise background averaged epistatic terms. The type of epistatic motif (categories) between mutations at a pair of positions determines whether these mutations can be incorporated by an evolutionary process proceeding by single mutation steps ^28,45^. Of the four categories, sign epistasis and reciprocal sign epistasis are the extreme forms that limit the accessible trajectories; their prevalence is a direct measure of ruggedness of the fitness landscape ^46,47^. Shown here are the frequencies of each epistatic motif amongst all significant pairwise terms, indicating substantial extreme epistasis.

**Extended Data Figure 7:**
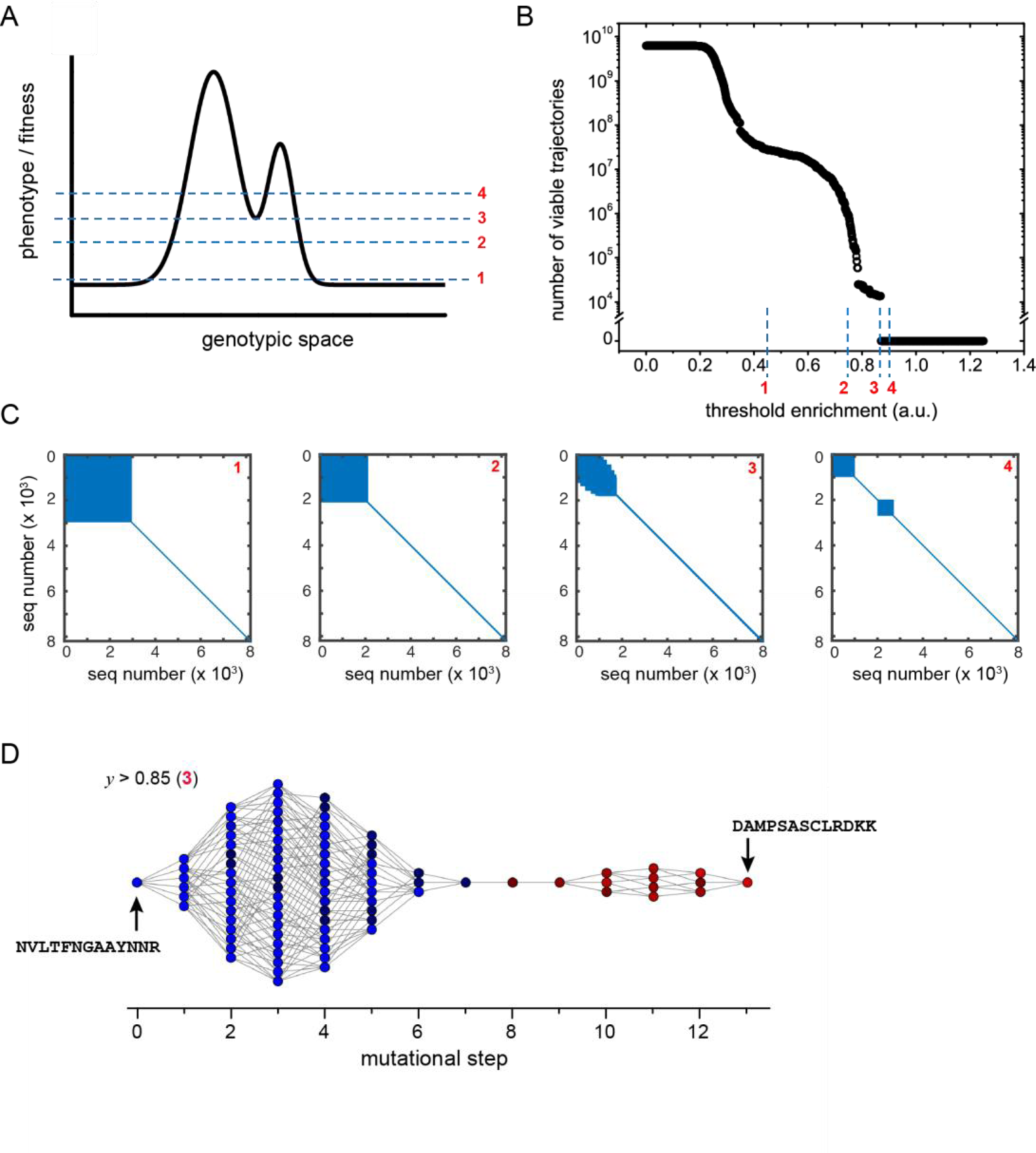
Functional connectivity of the sequence space. **A**), A schematic illustrating the concept of genotypic connectivity as a function of different phenotypic thresholds (marked in red, 1-4). The solution space is connected up to threshold 3. (**B**) The number of single-step trajectories as a function of threshold brightness for the data; thresholds corresponding to the cartoon in panel A are indicated. Note that the solution space becomes disconnected (zero viable paths) at threshold 3. (**C**) Genotypic “connectograms”, a graphical representation of the single step functional connectivity of the sequence space as a function of threshold (red numbers corresponding to panel B) (see Methods for computational process). The graphs show that threshold 3 represents the critical point after which the solution space breaks and is not fully connected. (**D**), The structure of the solution space at the threshold for functional connectivity. As in Figure 5, the space is dumbbell shaped, but with the neck linking the space now defined by an ordered series of single mutants. These genotypes (steps 7-9) harbor a reciprocal sign epistatic motif, which defines the strict order in which these mutations must come in order to be functionally connected. Note that at this threshold (y=0.85), the starting and ending genotypes shown here are not the parental ones.

